# Integrating single-cell RNA-Seq with spatial transcriptomics in pancreatic ductal adenocarcinoma using multimodal intersection analysis

**DOI:** 10.1101/254375

**Authors:** Reuben Moncada, Florian Wagner, Marta Chiodin, Joseph C. Devlin, Maayan Baron, Cristina H. Hajdu, Diane M. Simeone, Itai Yanai

**Author notes:** Materials & Correspondence.

## Abstract

To understand tissue architecture, it is necessary to understand both which cell types are present and the physical relationships among them. Single-cell RNA-Seq (scRNA-Seq) has made significant progress towards the unbiased and systematic identification of cell populations within a tissue, however, the characterization of their spatial organization within it has been more elusive. The recently introduced ‘spatial transcriptomics’ method (ST) reveals the spatial pattern of gene expression within a tissue section at a resolution of a thousand 100 µm spots across the tissue, each capturing the transcriptomes of multiple cells. Here, we present an approach for the integration of scRNA-Seq and ST data generated from the same sample, and deploy it on primary pancreatic tumors from two patients. Applying our multimodal intersection analysis (MIA), we annotated the distinct micro-environment of each cell type identified by scRNA-Seq. We further found that subpopulations of ductal cells, macrophages, dendritic cells, and cancer cells have spatially restricted localizations across the tissue, as well as distinct co-enrichments with other cell types. Our mapping approach provides an efficient framework for the integration of the scRNA-Seq-defined subpopulation structure and the ST-defined tissue architecture in any tissue.

## INTRODUCTION

Recent technological advances have enabled a view into cancer at unprecedented molecular resolution^1^. Single-cell RNA-sequencing (scRNA-Seq) has emerged as a powerful tool for the unbiased and systematic characterization of the cells present in a given tissue^2–4^. Indeed, the application of scRNA-Seq to patient tumors has uncovered multiple cellular subpopulations and has highlighted intercellular cross-talk within the tumor microenvironment^5–12^. However, due to the necessity of cellular dissociation prior to sequencing of individual cells, the spatial context for each cell is lost, thus limiting insight into the precise spatial organization of the cell types constituting a tumor.

Methods providing spatially resolved transcriptomic profiling^13–19^ have also been introduced, and these are useful for the integration with scRNA-Seq data. For example, *in situ* hybridization (ISH) gene expression atlases serve as useful references for cellular localization^20,21^. Using the ISH atlas as a guide, it is possible to accurately map rare subpopulations using a small subset of genes. Such atlases do not exist for solid tumors though, which have variable tissue architecture and gene expression patterns. Thus, high-throughput and comprehensive mapping of single-cells onto tissue requires robust integration of multiple methods.

The spatial transcriptomics (ST) method enables spatially-resolved transcriptomic profiling of tissue sections using spatially barcoded oligo-deoxythymidine (oligo-dT) microarrays, allowing for unbiased mapping of transcripts over entire tissue sections^22^ (Fig. 1a). The ST method has already been used to study the mouse olfactory bulb^22^, breast cancer^22^, melanoma^23^, prostate cancer^24^, gingival tissue^25^, adult human heart tissue^26^, mouse and human spinal cord tissue^27^, as well as different plant species^28^. Similarly to the other previously reported spatially-resolved transcriptomic tools^13,29,30^ however, a main limitation of the ST method is its lack of cellular resolution; each spot captures the transcriptomes of as few as 10 to as many as 200 neighboring cells, depending on the tissue context^31^. While scRNA-Seq and ST data each have limitations, we reasoned that these could be overcome by an integration of the two data modalities to enable comprehensive and unbiased tissue analysis.

**Figure 1.**
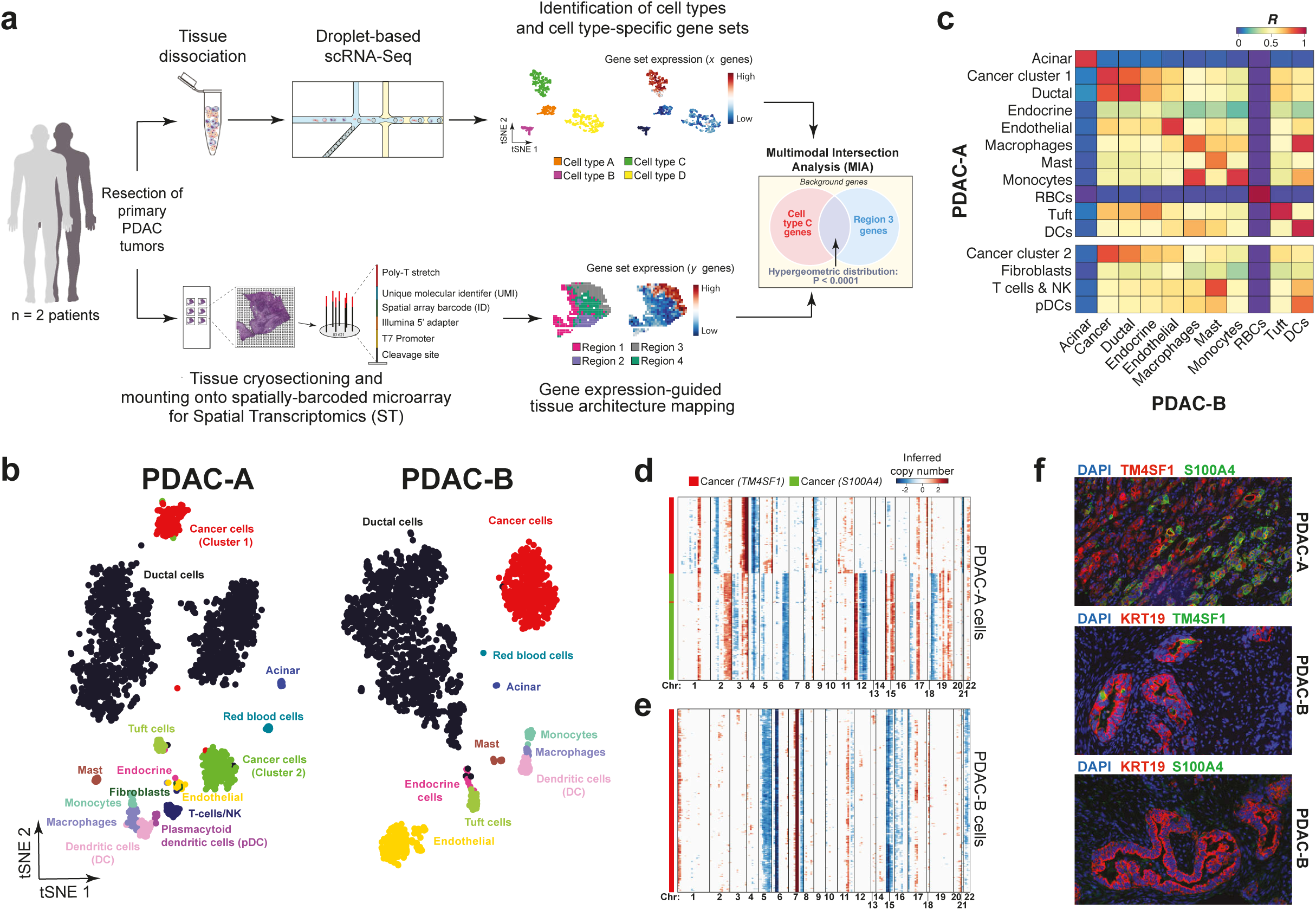
scRNA-Seq analysis of two PDAC patient tumors. (a) Schematic of the experimental design and analysis. Surgically resected PDAC tumors are split and processed in parallel for scRNA-Seq and ST. After clustering, the cell type of each cluster is inferred according to specifically expressed genes. Cryosections of the rest of the sample was used for ST analysis in which each spot captures the transcriptomes of the cells at a specific location in the tissue. Applying our Multimodal Intersection Analysis (MIA) across the two datasets reveals the spatial distribution of the cell populations and subpopulations. (b) t-SNE projection of 1,926 cells from the PDAC-A tumor (left panel) and 1,733 cells from PDAC-B (right panel). Clusters are colored and labeled according to their inferred cell type identities. (c) Correspondence between PDAC-A and PDAC-B cell types computed by Pearson’s correlation on the average cell type transcriptome. Shared cell types are shown in the top panel and cell types identified only in PDAC-A are indicated in the bottom panel. (d-e) CNV profiles inferred from scRNA-Seq on PDAC-A (d) and PDAC-B (e). Red and blue indicate chromosomal amplifications and deletions, respectively. (f) Double immunofluorescence staining of markers for cancer cell subpopulations. Top panel, double staining of TM4SF1 (cancer cluster 1) and S100A4 (cancer cluster 2) in PDAC-A formalin-fixed paraffin embedded (FFPE) tissue. Note the mutual exclusion of TM4SF1 and S100A4 signals. Middle, KRT19 and TM4SF1 staining in PDAC-B tissue. Bottom, KRT19 and S100A4 staining in PDAC-B tissue. Note colocalization of KRT19 and TM4SF1 signals, but a lack of S100A4 signal in PDAC-B as expected.

Here, we present an integration of scRNA-Seq with the ST method in pancreatic adenocarcinoma (PDAC). In this approach, a tumor is first bisected and a single-cell suspension is generated from one half and processed for scRNA-Seq to identify the cell populations present in the tissue. From the second half, cryosections are processed using the ST method to provide an unbiased map of expressed transcripts across the tissue. We then integrate these two datasets by introducing multimodal intersection analysis (MIA), in which we infer the enrichment of specific cell types in a given tissue region by assessing the degree of overlap between ST genes specific for that region and the cell type-specific gene expression as defined by the scRNA-Seq data. Studying two primary PDAC tumors from different patients we identified enrichments of specific cell types and subpopulations across spatially restricted regions of the tissue. Our approach for combining these two complementary and powerful technologies has the advantage that it is easily scalable to any architecturally complex tissue.

## RESULTS

### Identifying cell populations in pancreatic ductal adenocarcinoma with single-cell RNA-Seq

We processed fresh primary PDAC tumors from two untreated patients – henceforth, PDAC-A and PDAC-B – for parallel scRNA-Seq and ST analysis (Fig. 1a, see Methods). The scRNA-Seq data consisted of cells with approximately 2,500 to 3,300 unique molecular identifiers (UMIs) and approximately 1,400 to 1,700 uniquely expressed genes per cell (Fig. S1a-d). To infer cell type identities, we used a recursive hierarchical clustering scheme (see Methods) that applies our recently developed *k*-nearest neighbor (KNN) smoothing algorithm^32^ to reduce the noise inherent to scRNA-Seq data^33^. We identified fifteen and eleven distinct populations in the PDAC-A and PDAC-B tumors, respectively, providing an in-depth perspective of the PDAC tumor microenvironment (Fig. 1b). The shared cell types between the patient samples demonstrated strong correlation based on their gene expression profiles, suggesting consistent annotation of clusters across samples (Fig. 1c).

To distinguish the PDAC cancer cells from the non-malignant ductal cells and other epithelial cells, we used scRNA-Seq-based copy number variation (CNV) analysis (as performed previously^5^, see Methods). For each identified cell type, we inferred chromosomal amplifications and deletions (with the cells of other clusters as the background) and detected two clusters in PDAC-A and one cluster in PDAC-B that displayed aberrant CNV profiles (Fig. 1d-e). Notably, the chromosomal loss along chromosome 6 in both samples and the chromosomal gain on chromosome 7 in the PDAC-B profile is consistent with the most common chromosomal abnormalities seen in PDAC from cytogenetic data^34^. As a negative control, we removed a random subset of ductal cells from the background and inferred the relative CNV profiles of these cells (Fig. S2a) and found that these do not show an aberrant CNV profile (Fig. S2a). The two PDAC-A cancer clusters – henceforth, cancer clusters 1 and 2 – were enriched for expression of either *TM4SF1* (cluster 1) or *S100A4* (cluster 2), while the PDAC-B cancer cluster was only enriched for *TM4SF1* expression (Fig. S2b), suggesting similarity between PDAC-A cancer cluster 1 and the PDAC-B cancer cluster.

To further validate whether the *TM4SF1* and *S100A4* expressing populations identified in the scRNA-Seq data represent cancer cell populations, we performed double immunofluorescence (IF) staining of TM4SF1 and S100A4 on formalin-fixed paraffin embedded (FFPE) tissue originating from the same patients (Fig. 1f, Fig. S2c-d). We found TM4SF1 and S100A4 signals in malignant PDAC-A ducts (Fig. S2c-d, top panels) but not in non-malignant ducts (bottom panels). Interestingly, the PDAC-A cancer cells displayed mutually exclusive staining for TM4SF1 and S100A4 (Fig. 1f, top). When we double-stained PDAC-B tissue for KRT19 and TM4SF1 or KRT19 and S100A4, we found co-localization of the KRT19 and TM4SF1 signal (Fig. 1f, middle) but no co-localization of the KRT19 and S100A4 signals (Fig. 1f, bottom). Together with the distinct CNV profiles for the cancer clusters, the IF signals from the scRNA-Seq identified markers support the delineation of multiple distinct cancer cell populations.

### Spatial transcriptomics of PDAC tissue

To generate unbiased transcriptomic maps of the tissue sections, we mounted cryosections of the two unfixed PDAC tissues originating from the same tumor sample onto the spatially barcoded ST microarray slides (see Methods). After staining the tissue with hematoxylin and eosin (H&E), the slide was presented to a pathologist (C.H.H.) for histological annotation (Fig. 2a-b). In the PDAC-A tumor section, we defined four main regions: cancer cell ducts and desmoplasia, non-malignant duct epithelium, stroma, and normal acini-rich pancreatic tissue (Fig. 2a and Fig. S3a-d). The PDAC-B tissue section (Fig. 2b) did not appear to contain normal pancreatic tissue, however we noted the presence of interstitial space adjacent to the cancer (Fig. S3e-f).

**Figure 2.**
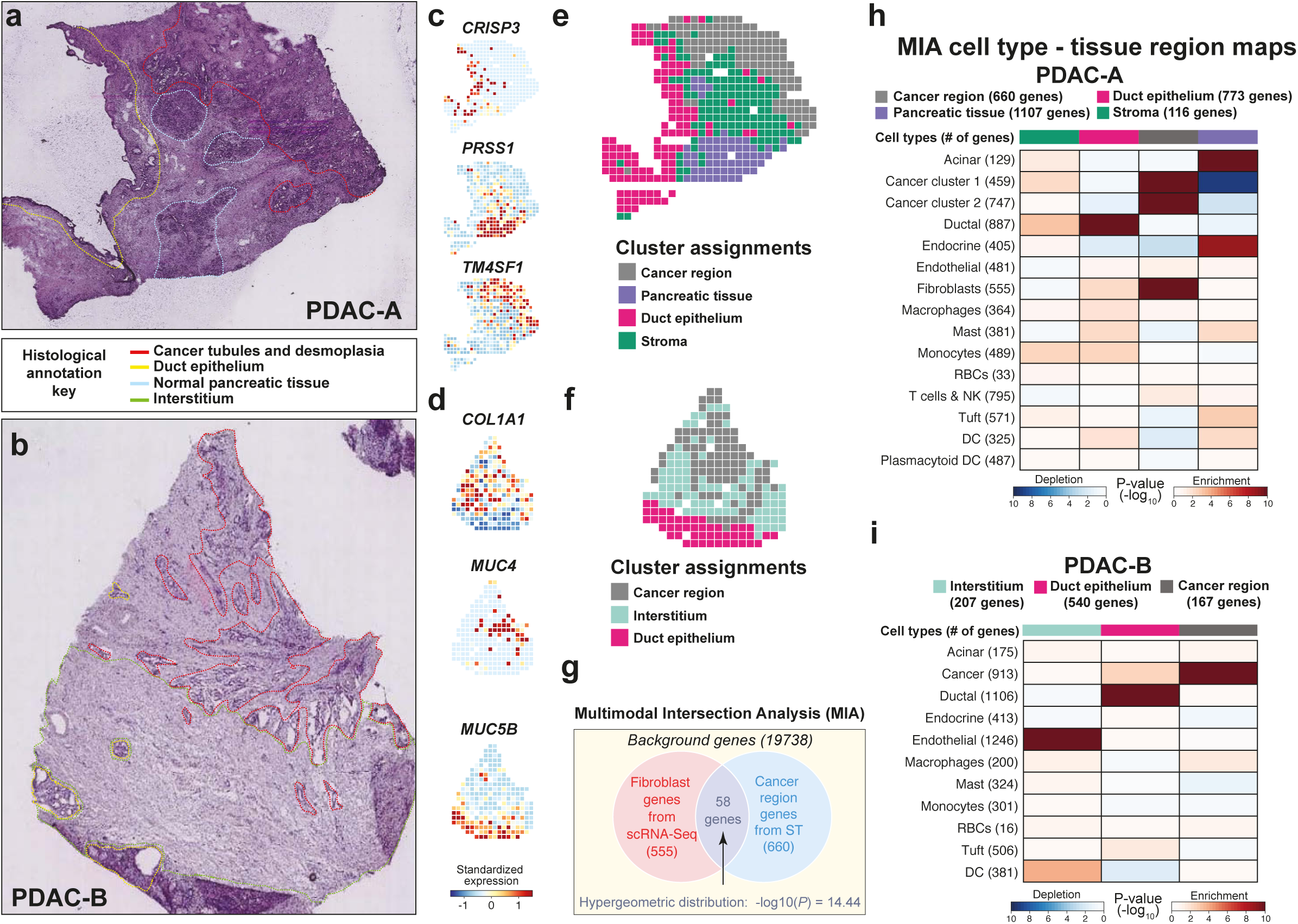
Spatial transcriptomics of PDAC and mapping of cell types. (a) Annotated PDAC-A tumor cryosection on the ST slide. The annotations indicate a region high in cancer cells and desmoplasia (red), normal pancreatic tissue (blue), and normal duct tissue (yellow). (b) Annotated PDAC-B tumor cryosection on the ST slide. Annotated regions include a cancer cell rich region (red), ducts (yellow), and interstitium (green). (c-d) Standardized expression levels of three genes in the PDAC-A ST (c) and PDAC-B ST (d) dataset. (e-f) Clustering of the PDAC-A and PDAC-B ST spots. Color indicates the clustering assignments. (g) Multimodal intersection analysis (MIA). The hypergeometric distribution is used to infer the significance of the intersection of genes enriched in a given cell type (fibroblasts in this case) and genes enriched in a given tissue region (cancer region). Applying this analysis systematically for all pairs of cell types and tissue regions allows for insight into the spatial distribution of the cell types in the tumor. (h) The PDAC-A MIA map of all scRNA-Seq identified cell types and ST defined regions. Each element in the matrix is computed as described in (d). Red indicates enrichment (significant overlap), blue indicates depletion (significantly low overlap). The bar on top indicates the regions delineated in (c). (i) The PDAC-B MIA map.

The samples were then processed for ST analysis, involving cDNA synthesis, amplification by *in vitro* transcription, library construction, and sequencing^19^. We demultiplexed the sequenced reads and identified their spatial location within the tissue using the ST location-specific barcodes of the array. By our estimation from the H&E images, the ST spots captured approximately 20-70 cells per spot (Fig. S4). We expect these numbers to be highly variable depending on the type of tissue and location within the tissue queried. For example, highly fibrotic tissue regions enriched in connective tissue (therefore, lower cellular content) may not capture many cells in ST, compared to tissue with high cellular density such as the acinus of the pancreas (Fig. S4). We detected approximately 2,400 UMIs and approximately 1,000 unique genes per spot for both ST datasets (Fig. S5a-f). Other published works using the ST method (particularly on human tissue) report similar statistics for the average number of UMIs per spot (Table S1). In both sample datasets, we found that the spatial expression of many dynamically expressed genes matched the annotated histological regions (Fig. 2c-d).

### Multimodal Intersection Analysis

To integrate the scRNA-Seq and ST datasets, we developed Multimodal Intersection Analysis (MIA). This analysis proceeds by first delineating sets of cell type-specific and tissue region-specific genes and then comparing their patterns of enriched or depleted intersections. In the scRNA-Seq data, we defined the gene sets by identifying for each cell type those genes whose expression is statistically higher in the cells annotated to that cell type in comparison to expression in the remaining cells (*P* < 10^-5^, two-tailed Student’s t-test, see Methods). In parallel, in the ST data we defined sets of genes with specific expression for each spatial region. For this, we first performed principal components analysis (PCA) on the 200 most dynamically expressed genes across all ST spots (Fig. S5g-h, see Methods). After clustering the spots based on the PC scores (see Methods), we found that the clusters consistently coincided with tissue histological annotations (Fig. 2e-f and Fig. S5g-h). We then identified genes whose expression is significantly enriched for expression in each region (*P* < 0.01, two-tailed Student’s t-test).

With the gene sets extracted across the scRNA-Seq and ST modalities, MIA next computes the hypergeometric distribution in order to assess the overlap between each pair of cell type-specific and region-specific gene sets (Fig. 2g). As an example, we found that the set of genes enriched for expression in the fibroblasts of PDAC-A overlaps significantly with the set of genes with enriched expression in the cancer region of the ST data (Fig. 2h, *P* < 10^-10^). Thus, while the scRNA-Seq data led to identification of fibroblasts, MIA further revealed that these were enriched in the cancer region, as opposed to the stromal region. This suggests that the transcriptomes identified for the fibroblasts correspond to that of activated fibroblasts^35^ (Fig. 2h). Extending this analysis to all pairs of cell types and tumor regions, we found that the duct epithelium region was mainly enriched with ductal cells, while the pancreatic tissue was enriched with acinar cells and endocrine cells (*P* < 10^-10^, for all specified enrichments). A similar MIA map was found for the PDAC-B sample (Fig. 2i). Interestingly, in PDAC-B the interstitium was enriched for endothelial cells and dendritic cells. These results support the utility of MIA maps to provide spatial and functional annotations for the scRNA-Seq-defined cell populations.

### Identification and mapping of cell type subpopulations across tissue regions

One of the most useful aspects of scRNA-Seq is its ability to distinguish distinct subpopulations within cell types. We thus sought to characterize the intra-population heterogeneity by identifying subpopulations within cell types and then apply MIA to ask if they map in a spatially-restricted manner across the tumor tissue. From both tumor samples, a majority of the scRNA-Seq cells consisted of *KRT19* expressing ductal cells (Fig. 1b, S2b), one of the two main constituent cell types of the pancreatic exocrine system^36^. We identified a total of four ductal subpopulations: a ductal population expressing *APOL1* and hypoxia-response related genes including ERO1A^37^ and *CA9*^38^, a terminal ductal population expressing *TFF1, TFF2,* and *TFF3*^39^, a centroacinar ductal population expressing *CRISP3* and *CFTR*^39^, and antigen-presenting ductal cells expressing major histocompatibility complex (MHC) class II genes *CD74, HLA-DPA1, HLA-DQA2, HLA-DRA, HLA-DRB1, HLA-DRB5,* and complement pathway components *C1S, C4A, C4B, CFB,* and *CFH* (Fig. 3a-d). Although MHC class II molecules are primarily expressed on the surface of professional antigen-presenting cells (B-cells, macrophages, dendritic cells), epithelial cells in the liver, gastrointestinal and respiratory tracts are known to express MHC class II^40,41^. Because of their presence in the tumor, it is likely that these non-professional antigen-presenting ductal cells play a role in modulating the inflammatory response within the microenvironment by promoting T cell activation^40,41^. When we performed double IF of archival FFPE patient tissue, we found co-localization of subpopulation markers with the duct marker KRT19, confirming the presence of these ductal cell subpopulations (Fig. 3e-h, Fig. S6).

**Figure 3.**
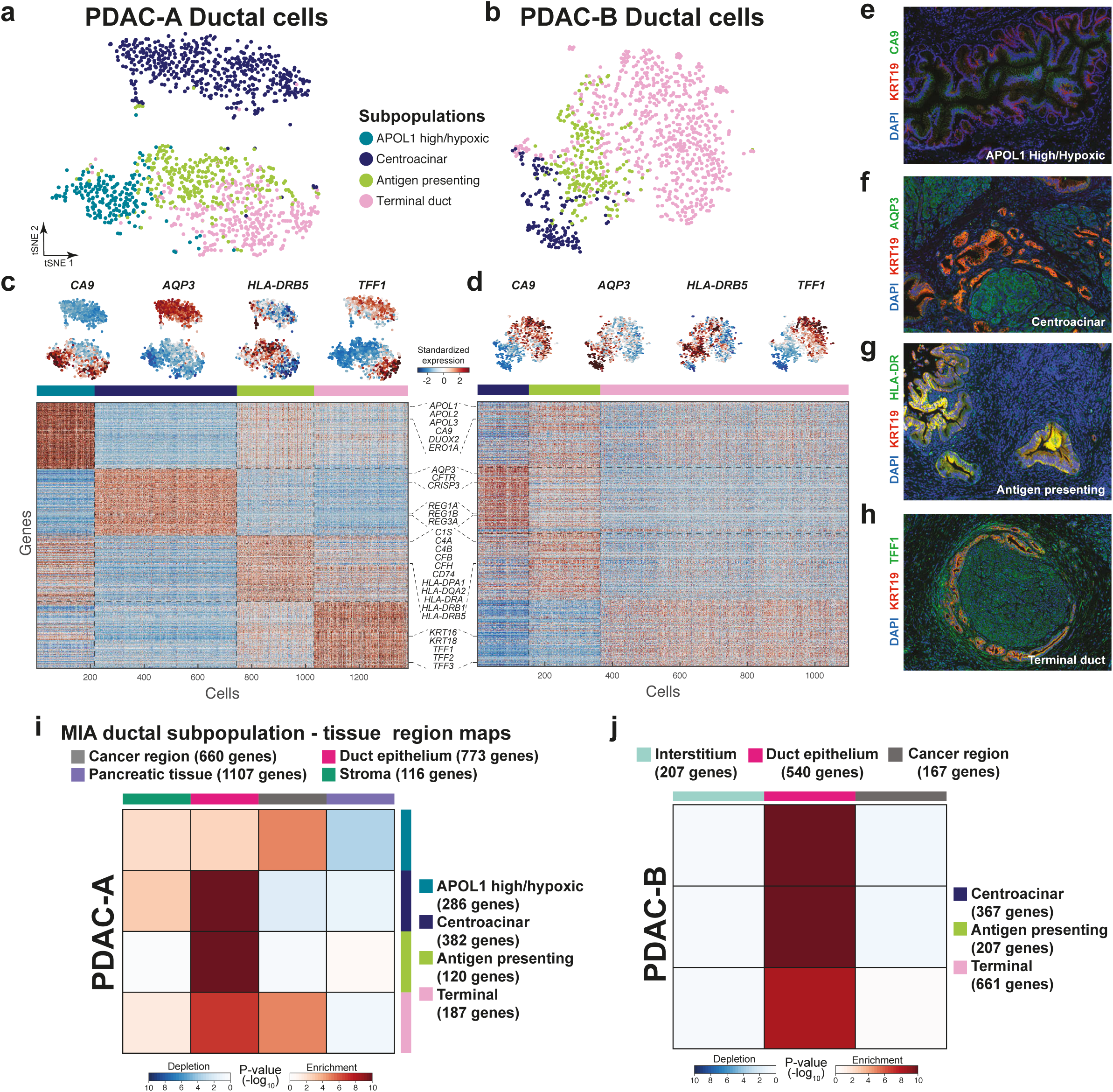
MIA mapping of ductal subpopulations across tissue regions. (a-b) Identifying subpopulations of ductal cells in PDAC-A (a) and PDAC-B (b). Colors in these t-SNE projections indicates the identified subpopulations. (c-d) The expression levels are shown for the indicated genes with subpopulation-specific patterns. The heatmaps show the standardized expression of the top 200 genes with subpopulation-specific expression. The genes are ordered identically across the PDAC-A and PDAB-B heatmaps. (e-h) Double immunofluorescence staining of KRT19 (ducts) and subpopulation markers CA9 (e, APOL1 high/hypoxic ductal cells), AQP3 (f, centroacinar ductal cells), HLA-DR (antigen presenting ductal cells), and TFF1 (h, terminal ductal cells). (i-j) MIA maps of PDAC ductal subpopulations across the ST regions identified in Figure 2.

As in our cell type analysis, we identified marker genes specific to each ductal subpopulation and determined the enrichment of these ductal subpopulations across the tissue region using MIA. We found that all ductal subpopulations in PDAC-A were enriched in the duct region of the tissue while only the hypoxic and terminal ductal cell populations were significantly enriched in the cancer region (*P* < 10^-4^, Fig. 3i). It is possible that the transcriptional phenotypes exhibited by these ductal cells reflect environmental signals from the surrounding tissue; *i.e.*, hypoxic ductal cells may be most enriched in the cancerous region compared to the ducts because this region is more hypoxic than other regions. Additionally, the hypoxic ductal cells also appeared to be depleted in the pancreatic tissue (*P* < 10^-3^, Fig. 3i). In contrast, the ductal subpopulations in PDAC-B were all exclusively enriched in the ducts of the tissue (*P* < 10^-4^, Fig. 3j).

We also found that the PDAC-A macrophages comprised two subpopulations, one of which expressed *IL1B,* corresponding to an inflammatory M1 state,^42,43^ and the other resembling the M2 alternatively activated state based on its expression of *CD163* and *MS4A4A*^44,45^ (Fig. S7a). We also found two subpopulations of dendritic cells, one of which expressed higher levels of complement pathway genes and MHC class II (Fig. S7b). When we tested for the enrichment of these subpopulations across the tissue with MIA, we found that the subpopulations of macrophages and dendritic cells appear to have opposite patterns of enrichment across the tissue (Fig. S7b,d). The M2-like macrophages appeared most enriched in the ducts while the M1 macrophages are more enriched in the stroma and cancer regions (Fig. S7c). This result is consistent with the M2-like population being tissue resident, with its location reflecting its endogenous functional location, while the M1-like population localizes specifically to the cancer and stroma regions. For the dendritic cell subpopulations, one subpopulation appeared most enriched in pancreatic tissue, while a second population appeared most enriched in the ducts of the tissue (Fig. S7d). Based on the MIA maps, it is likely that these subpopulations play unique roles in the tissue based on their differential localization thus highlighting new research directions.

### Mapping of distinct cancer populations across tissue

In the scRNA-Seq data for the PDAC-A tumor we found two cancer cell populations that appeared to be both genetically and transcriptionally distinct (Fig. 1b-e). Although we found both cancer populations to be highly enriched in the corresponding ST cancer region (Fig. 2d), we asked whether the two cancer populations co-localized with different cell types within this region. To address this question, we used hierarchical clustering to divide the cancer region into transcriptionally coherent sub-regions (Fig. 4a-b). After defining sets of genes enriched for expression in each sub-region, we again applied our MIA mapping approach to study the overlap between the sub-regions sets and the sets of genes defined above for each cell type/subpopulation from the scRNA-Seq data.

**Figure 4.**
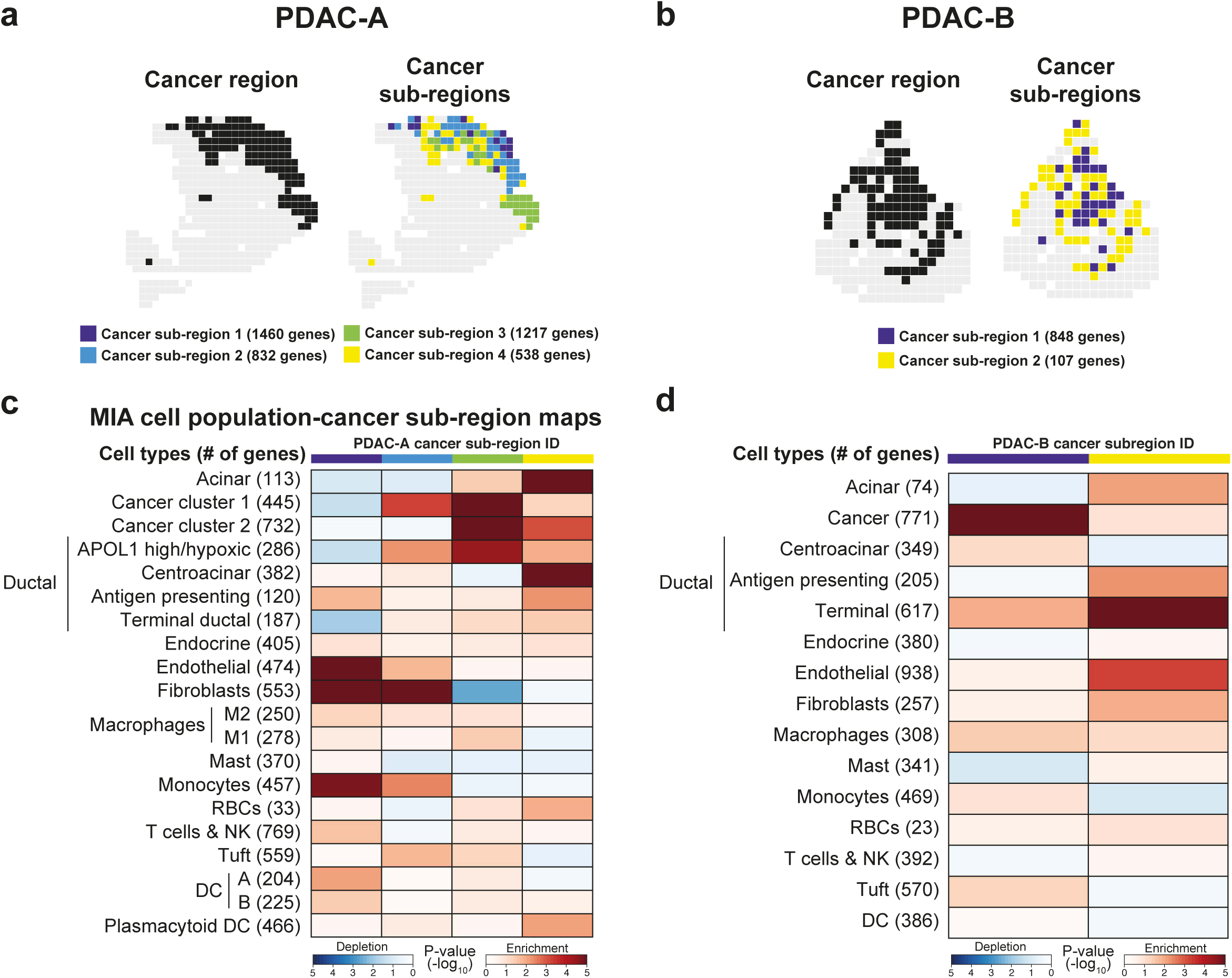
Cancer sub-regions reveal differential cell type and subpopulation enrichments. (a-b) Identifying sub-regions in the PDAC-A (a) and PDAC-B (b) ST cancer regions. The colors indicate the cluster assignments exclusively for the spots annotated to be in the cancer regions identified in Figure 2. (c-d) MIA maps of the cancer identified sub-regions in PDAC-A (c) and PDAC-B (d) and the cell type and cell subpopulations identified in Figure 1 and 3.

The PDAC-A sub-region 1 exhibited enrichments of monocytes, endothelial cells and fibroblasts, but a relative depletion of cancer cells. In the second sub-region, we found enrichment of only the cancer cluster 1 subpopulation together with fibroblasts, suggesting a potential specific relationship for one of the cancer subpopulations. The third sub-region revealed an enrichment of both cancer subpopulations with hypoxic ductal cells and an absence of fibroblasts, while the last sub-region revealed co-enrichment of the acinar cells, cancer cluster 2, antigen presenting and centroacinar ductal cells, and plasmacytoid dendritic cells (Fig. 4c, *P* < 0.01 for all enrichments).

In the PDAC-B sample, we characterized two sub-regions within the cancer region and defined a gene set specific to each. In the first sub-region, we found co-enrichment of cancer cells with terminal ductal cells and macrophages, while the second sub-region was enriched for terminal ducts, fibroblasts, and endothelial cells (Fig. 4d). Across the two MIA maps, we find consistent mutual exclusion of antigen-presenting ductal cells and the *TM4SF1* expressing cancer populations (cancer cluster 1 for PDAC-A), suggesting a possible antagonistic relationship between the immunosuppressive cancer cells and antigen presenting cells. For example, the cancer cells may suppress antigen presentation but cells in the adjacent areas may be able to elicit an inflammatory response functioning as a containment method. We also observed consistent co-enrichment of fibroblasts with endothelial cells across the MIA maps, highlighting a potential relationship between the establishment of a fibrotic environment (by activated fibroblasts) and the formation of vasculature (by endothelial cells). Collectively, these results demonstrate the utility of MIA maps for generating hypotheses – requiring further testing – on the relationships of cell populations with spatially restricted architectures, even within a seemingly uniform histological feature.

## DISCUSSION

We have presented here a method for the identification and spatial mapping of distinct cell types together with their distinct subpopulations within heterogeneous samples. The method begins with the characterizing of cell types and subpopulations present in a tissue by scRNA-Seq, and in parallel, the identifying of transcriptomic regions from a portion of the same tissue using by spatial transcriptomics (ST). We then take advantage of the unbiased nature of both data modalities to detect enrichment of cell populations across the spatial regions using our multimodal intersection analysis (MIA). By applying this approach to two PDAC samples we mapped relationships among the distinct and reproducible cell populations intrinsic to the tumor microenvironment (immune, stromal, and malignant cells) and those comprising the normal pancreatic tissue (pancreatic ductal cells, acinar cells, endocrine cells). We demonstrated that MIA can be used to infer the state of a cell population given its location (see activated fibroblasts in Figure 2), the distinct localizations of sub-populations of a cell type (ductal subpopulations in Figure 3), and that relationships among the cell types can be identified with in sub-regions of the tumor (Figure 4).

Established methods for mapping transcripts (ISH, FISH) or proteins (immunohistochemistry, IHC) are limited to a few antibodies or *in situ* probes per tissue section. Multiplexing of these also leads to a limitation in the number of targets to visualize simultaneously. The ST method has the advantage in this respect in that it provides an unbiased map of expressed transcripts in a given tissue section^24–26,28,46^. Although other spatially resolved transcriptomic methods can detect hundreds of RNA species at single cell resolution^13^, there remains a limit on the depth of information and they cannot, for example, identify all genes specifically enriched for expression to a defined spatial region of the tissue. ST is thus uniquely positioned for studying histologically variable tissues, such as tumors, without ISH maps to guide cell location inference^20,21^. Instead of querying specific markers using ISH to characterize the scRNA-Seq results, the integration ST enables a more efficient initial interrogation of a tissue.

Our approach to integrating scRNA-Seq and ST using MIA has the advantage that the two modalities enable a better understanding of cell populations and subpopulation structure. Because the ST spots are clustered from the transcriptomic data independent of tissue histology, MIA maps provide an unbiased perspective into the organization of distinct populations in local tissue niches. This allows for the inference of functional relationships between scRNA-Seq-defined populations based on their co-localization in space, and ultimately provides a more comprehensive characterization of cell types in their native environment that cannot be gained from either modality alone. The unique perspective offered by the intersection of these two technologies is likely to advance clinical/pathological investigation of tumor tissue beyond typical histological techniques. We imagine that the modern pathologist will be able to move beyond studying cell type-specific markers and use the maps created by our approach to perform more precise diagnoses of the cancer stage, type, and history.

Despite the potential of this integrated method, there are a number of limitations. First, the current version of the ST array is about 6 x 7 mm in size, consisting of just over 1000 spots, with each spot being 100 µm in diameter and 100 µm apart. Thus, the array is neither large enough to cover the entire tissue (in many cases) nor are the spots small enough to provide single-cell resolution. We expect the ST technology to improve in terms of both the number and size of the spots on the array, as well as the technical protocol which would reduce the diffusion of transcripts across spots as much as possible. The process of tissue dissociation necessary for scRNA-Seq has the potential to lead to biases as some cell types may be more amenable to dissociation over others, leading to overrepresentation in the data. Therefore, it is necessary to thoroughly dissociate the tissue for processing in order to gain a full representation of the cell types present in the tissue. Additionally, the tissue dissociated for scRNA-Seq comes from the same biological sample, but is not the same exact tissue used for ST. For reliable mapping of scRNA-Seq identified cell types in the ST data it is crucial for the scRNA-Seq to capture the most abundant cell types present in the tissue. Despite these limitations, ST remains very accessible to researchers as it requires only a few additional steps compared to bulk RNA-Seq analysis of homogenized tissue; unlike other spatially resolved transcriptomic approaches^13,14^, these additional steps do not require specialized equipment outside of what is already found in most laboratories or research institutions^22^.

Analyzing the same tumor sample with multiple modalities will likely have far-reaching implications, particularly with regard to the identification and classification of cell populations that constitute a heterogeneous tissue. The advent of scRNA-Seq has allowed for the identification of cancer subpopulations and non-malignant cell subpopulations^10,47^; thus MIA mapping can aid in assigning potential functional roles of cellular subpopulations based on spatial localization (relative to the tissue, or relative to the other cells present). By applying scRNA-Seq and ST on the same biological sample as we describe here, rare subpopulations specific to the sample can be mapped to the same tissue of origin. Moreover, therapeutically targetable cell populations identified by our approach can be better understood by being mapped to their relative environments in the tissue, as well as by mapping the other cell types that are co-localized with them. In the case of tumors for which the precise composition of different tumor sub-classifications is likely to vary from individual to individual, the subpopulation composition and spatial localization can be ascertained for a given patient and can perhaps be correlated with patient outcome.

## METHODS

### Tumor sample handling and dissociation to a single-cell suspension

Two pancreatic ductal adenocarcinoma tumors were delivered in RPMI (Fisher Scientific) on ice directly from the operating room to the lab after clearing pathology (∼2 hours). Each tumor resection was rinsed in ice cold PBS and cut into ∼4-5 mm^3^ pieces from which 1 mm thick slices were taken and set aside in ice-cold PBS. The remaining ∼3-4 mm^3^ pieces were embedded in chilled OCT and snap-frozen in isopentane cooled with liquid N_2_. The 1 mm tissue slices stored in PBS was further minced with scalpels to < 1 mm^3^. Tissue was rinsed from the dish with ice cold PBS and pelleted by centrifuging at 300 x g for 3 minutes at 4 degrees. PBS was aspirated and 5 ml 0.25% pre-warmed trypsin-EDTA with 10 U/µl DNaseI (Roche) was added and put into a 37°C water bath for 30 minutes with gentle inversion every 5 minutes. The resulting suspension was filtered through a 100 µm cell strainer to remove larger chunks of undigested tissue. Enzymatic digestion was quenched with the addition of FBS to a final concentration of 10%. Cells were pelleted by centrifuging the suspension at 300 x g for 3 minutes at 4 degrees and washed twice with 5 ml ice-cold PBS. After a final spin at 300 x g for 3 minutes, the cells were resuspended in PBS to a final concentration of 10,000 cells/ml. The resulting viability was >95% as shown by trypan blue exclusion.

### inDrop library preparation and scRNA-Seq

From each single-cell suspension, between 6,000 and 12,000 cells were encapsulated using the inDrop platform and reverse transcription (RT) reaction was performed as previously described^39^. The number of PCR cycles performed for final library amplification ranged from 9-12 cycles. Libraries were diluted to 4 nM and paired end sequencing was performed on an Illumina NextSeq platform. Between 139 million and 145 million paired reads were generated for each library, corresponding to approximately 35,000 paired reads per cell.

### Processing of inDrop single-cell RNA-Seq sequencing data

Raw sequencing data obtained from the inDrop method were processed using a custom-built pipeline, available online (https://github.com/flo-compbio/singlecell). Briefly, the “W1” adapter sequence of the inDrop RT primer was located in the barcode read (the second read of each fragment), by comparing the 22-mer sequences starting at positions 9-12 of the read with the known W1 sequence (“GAGTGATTGCTTGTGACGCCTT”), allowing at most two mismatches. Reads for which the W1 sequence could not be located in this way were discarded. The start position of the W1 sequence was then used to infer the length of the first part of the inDrop cell barcode in each read, which can range from 8-11 bp, as well as the start position of the second part of the inDrop cell barcode, which always consists of 8 bp. Cell barcode sequences were mapped to the known list of 384 barcode sequences for each read, allowing at most one mismatch. The resulting barcode combination was used to identify the cell from which the fragment originated. Finally, the UMI sequence was extracted, and reads with low-confidence base calls for the six bases comprising the UMI sequence (minimum PHRED score less than 20) were discarded. The reads containing the mRNA sequence (the first read of each fragment) were mapped by STAR 2.5.1 with parameter “—outSAMmultNmax 1” and default settings otherwise^48^. Mapped reads were split according to their cell barcode and assigned to genes by testing for overlap with exons of protein-coding genes. Only single-cell transcriptomes with ≥ 1000 UMIs, ≤ 20% mitochondrial transcripts and ≤ 30% ribosomal transcripts were kept, leaving 1926 cells for the PDAC-A dataset, and 1733 cells for the PDAC-B dataset. UMI counts were normalized by the total number of transcripts per cell, and a scale factor equivalent to the median number of transcripts across all cells was applied (transcripts per median, TPM). Expression was transformed using Freeman-Tukey transform as described previously^32^. Mitochondrial and ribosomal genes were removed from the expression matrix for further analysis.

### K-Nearest Neighbors (KNN) smoothing of scRNA-Seq data

To reduce the noise inherent to scRNA-Seq data^49^, we applied our recently proposed KNN-smoothing method^32^. Briefly, all single-cell expression profiles were normalized to median number of total transcripts per cell, the Freeman-Tukey transformation was applied to all expression values, and the *k* closest neighbors of each cell were identified using Euclidean distance. The expression profile of each cell was then combined with those of its neighbors, thus obtaining its smoothed expression profile.

### Hierarchical clustering of single-cell RNA-Seq data and marker gene selection

For clustering and identification of cell types, we used a recursive clustering scheme involving data, smoothing, clustering, removal of identified clusters from the rest of the data, and repeated until all clusters are identified. Briefly, the data was first smoothed using our recently developed KNN-smoothing algorithm^32^, normalized by the total number of transcripts for that cell and scaled by a factor equal to the median transcript number across all cells, and transformed using the Freeman-Tukey transformation y = sqrt(x) + sqrt(x+1) for variance stabilization. Cells were then clustered with Ward’s criterion using the most variable genes (defined as Fano factor and mean expression above mean-dependent threshold). From resulting clusters, we obtained a list of marker genes by examining genes that are differentially expressed (*P* < 10^-5^, two-tailed Student’s t-test; effect size > 0.2, Cohen’s d). Once larger clusters were identified and removed from the data, the process was repeated and smoothed with lower values of *k* to prevent smoothing over of smaller clusters of cells until all clusters were identified. In this process, the annotated macrophages and dendritic cell clusters partitioned to two sub-clusters. We therefore kept these sub-clusters separate as they likely represented subpopulations. While clustering the PDAC-B data, we identified a population of ductal cells with low UMI counts and high mitochondrial content. Because of these reasons, and the lack of a similar population in PDAC-A, we suspected these cells to be low-quality cells that may have arisen as an artifact of the dissociation process. These cells were therefore removed from the PDAC-B dataset.

### Correlation between cell types in the scRNA-Seq data

To determine the similarity between cell types identified in PDAC-A and PDAC-B, the transcriptomes of all cells in each cluster were averaged (after TPM normalization and applying the Freeman-Tukey transform, see above), and the Pearson’s correlation coefficient was computed between the resulting averaged cell type profiles.

### Identification of ductal subpopulations

To sub-cluster the ductal subpopulations, the ductal cells were first isolated from the expression matrix and smoothed using KNN-smoothing^32^ with k = 64. Cells were then clustered with hierarchical clustering using the top variably expressed genes (see above). To annotate subpopulations, differentially expressed genes between each subpopulation was identified using a two-tailed Student’s t-test (selecting genes with *P* values less than 10^-5^). The top 200 differentially expressed genes for each subpopulation was used for the heatmap visualization shown in Figure 3c-d.

### Dimensionality reduction (PCA and tSNE)

Dimensionality reduction methods were performed on the Freeman-Tukey transformed data (after normalizing UMI counts to the median as described above) using variable genes (defined as Fano factor and mean expression above mean-dependent threshold). tSNE was performed using the following parameters: perplexity = 30 and initial dimension = number of principal components explaining >90% of the variance^50^.

### Cell cycle analysis of scRNA-Seq data

To perform the cell-cycle analysis shown in Figure S1, we used the ‘CellCycleScoring’ function included in the Seurat package v. 2.3^51^ to assign a cell-cycle phase to each cell in our scRNA-Seq datasets. List of genes specific for each cell-cycle phase are available at https://satijalab.org/seurat/cell_cycle_vignette.html.

### Copy Number Variation (CNV) analysis

CNV estimation was performed for each cell based on its expression profile using a similar approach to that previously described^5,7,9^. First, the PDAC-A and PDAC-B scRNA-Seq gene expression matrix was first smoothed with KNN-smoothing^32^, using k = 10. Next, genes were sorted based on their chromosomal location and a moving average of gene expression was calculated using a window size of 0.1 multiplied by the number of genes of each chromosome. The expression was then centered to zero by subtracting the mean. A subset of 200 randomly selected ductal cells were removed as a negative control for the analysis, leaving all remaining cells as the background.

### Selection of genes with population-specific expression patterns based on single-cell RNA-Seq data

To identify marker genes for each cell type or subpopulation, we identified differentially expressed genes as described above (*P* < 10^-5^, two-tailed Student’s t-test; effect size > 0.1, Cohen’s d). For each gene with *P <* 10^-5^, we examined for which cell type the effect size was highest and determined that gene to be specific for this cell type.

### Tissue preparation, cryosectioning, fixation, staining, and brightfield imaging for Spatial Transcriptomics (ST)

Patients at NYU Langone Health consented preoperatively to participate in the study. PDAC tumor tissue arrived in RPMI (Fisher Scientific) on ice. Tissue was gently washed with cold 1X-PBS, and 4-5 mm^3^ cubes were removed with a scalpel for OCT-embedding. Tissue was transferred from 1X PBS to a dry, sterile 10-cm dish and gently dried prior to equilibration in cold OCT for 2 minutes. The tissue was then transferred to a tissue-mold with OCT and snap-frozen in liquid nitrogen-chilled isopentane. Tissue blocks were stored at −80°C until further use. Prior to cryosectioning, the cryostat was cleaned with 100% ethanol, and equilibrated to an internal temperature of −18°C for 30 minutes. Once equilibrated, OCT embedded tissue blocks were mounted onto the chuck and equilibrated to the cryostat temperature for 15-20 minutes prior to trimming. ST slide was also placed inside cryostat to keep the slide cold and minimize RNase activity. Sections were cut at 10 µm sections and mounted onto the ST arrays, and stored at −80°C until use, maximum of two weeks. Prior to fixation and staining, the ST array was removed from the −80°C and into an RNase free biosafety hood for 5 minutes to bring to room temperature, followed by warming on a 37°C heat block for 1 minute. Tissue was fixed for 10 minutes with 3.6% formaldehyde in 1X PBS, and subsequently rinsed in 1x PBS. Next, the tissue was dehydrated isopropanol for 1 minute followed by staining with hematoxylin and eosin. Slides were mounted in 65 µl 80% glycerol and bright field images were taken on a Leica SCN400 F whole-slide scanner at 40X resolution.

### ST barcoded microarray slide information

Library preparation slides used were purchased from the Spatial Transcriptomics team (https://www.spatialtranscriptomics.com; lot 10002 and 10003). Each of the spots printed onto the array is 100 µm in diameter and 200 µm from the center-to-center, covering an area of 6.2 by 6.6 mm. Spots are printed with approximately 2 x 10^8^ oligonucleotides containing an 18-mer spatial barcode, a randomized 7-mer UMI, and a poly-20TVN transcript capture region^22^ (Fig. 1).

### On-slide tissue permeabilization, cDNA synthesis, probe release

After brightfield imaging, the ST slide was prewarmed to 42°C and attached to a pre-warmed microarray slide module to form reaction chambers for each tissue section. The sections were pre-permeabilized with 0.2 mg/ml BSA and 200 units of collagenase diluted in 1X HBSS buffer for 20 minutes at 37°C and washed with 100 µl 0.1X SSC buffer twice. Tissue was permeabilized with 0.1% pepsin in HCl for 4 minutes at 42°C and washed with 100 µl 0.1X SSC buffer twice. Reverse transcription (RT) was carried out overnight (∼18-20h) at 42°C by incubating permeabilized tissue with 75 µl cDNA synthesis mix containing 1X First strand buffer (Invitrogen), 5 mM DTT, 0.5 mM each dNTP, 0.2 µg/µl BSA, 50 ng/µl Actinomycin D, 1% DMSO, 20 U/µl Superscript III (Invitrogen) and 2U/µl RNaseOUT (Invitrogen). Prior to removal of probes, tissue was digested away from the slide by incubating the tissue with 1% 2-mercaptoethanol in RLT buffer (Qiagen) for one hour at 56°C with interval shaking. Tissue was rinsed gently with 100 µl 1X SSC, and further digested with proteinase K (Qiagen) diluted 1:8 in PKD buffer (Qiagen) at 56°C for 1 hour with interval shaking. Slides were rinsed in 2X SSC with 0.1% SDS, then 0.2X SSC, and finally in 0.1X SSC. Probes were released from the slide by incubating arrays with 65 µl cleavage mix (8.75 μM of each dNTP, 0.2 μg/μl BSA, 0.1 U/μl USER enzyme (New England Biolabs) and incubated at 37 °C for 2 hours with interval mixing. After incubation, 65 µl of cleaved probes was transferred to 0.2 ml low binding tubes and kept on ice.

### ST library preparation and sequencing

Libraries were prepared from cleaved probes as previously described, with the following changes: briefly, after RNA amplification by *in vitro* transcription (IVT) and subsequent bead clean-up, second RT reaction was performed using random hexamers, eliminating the need for a primer ligation step^52^. To determine the number of PCR cycles needed for indexing, 2 μl of the purified cDNA was mixed with 8 μl of a qPCR mixture [1.25x KAPA HiFi HotStart Readymix (KAPA Biosystems), 0.625 μM PCR lnPE1 primer, 12.5 nM PCR lnPE2 primer, 0.625 μM PCR Index primer, 1.25xEVA green (Biotium). Reactions were amplified on a Bio-Rad qPCR using the following program: 98 °C for 3 minutes, followed by 25 cycles of 98°C for 20 s, 60°C for 30?s and 72°C for 30 s. Optimal cycle number approximated to be the number of cycles required to reach saturation of signal minus 3-4 cycles to reach the exponential phase of the amplification. The remaining purified cDNA was indexed and using the same program described above, except amplified at the pre-determined number of cycles and with the inclusion of a final 5 minute extension at 72°C. Average lengths of the indexed, purified libraries were assessed using a 2100 Bioanalyzer (Agilent) and concentrations were measured using a Qubit dsDNA HS Assay Kit (Life Technologies), according to the manufacturer’s instructions. Libraries were diluted to 4 nM and paired-end sequencing was performed on an Illumina NextSeq sequencer with 31 cycles for read 1, and 55 cycles for read 2. Between 100 and 125 million raw read-pairs were generated for each sequenced library.

#### Primer sequences

*PCR InPE* 1:

5’-AATGATACGGCGACCACCGAGATCTACACTCTTTCCCTACACGACGCTCTTCCGATCT-3’

*PCR InPE 2*:

5’-GTGACTGGAAGTTCAGACGTGTGCTCTTCCGATCT-3’

*Cy3 anti-A probe:*

[Cy3]AGATCGGAAGAGCGTCGTGT

Cy3 anti-frame probe: [Cy3]GGTACAGAAGCGCGATAGCAG

### ST spot selection and image alignment

Upon removal of probes from ST slide, the slide is kept at 4°C for up to 3 days. The slide was placed into a microarray cassette and incubated with 70 µl of hybridization solution (0.2 µM Cy3-A-probe, 0.2 µM Cy3 Frame probe, in 1X PBS) for 10 minutes at room temperature. The slide was subsequently rinsed in 2X SSC with 0.1 % SDS for 10 minutes at 50°C, followed by 1 minute washes with 0.2X SSC and 0.1X SSC at room temperature. Fluorescent images were taken on a Hamamatsu NanoZoomer whole-slide fluorescence scanner. Brightfield images of the tissue and fluorescent images were manually aligned with Adobe Photoshop CS6 to identify the array spots beneath the tissue.

### ST library sequence alignment and annotation

Raw sequencing data obtained from the ST method were processed using a custom-built pipeline, available online (https://github.com/yanailab/celseq2). The pipeline was adapted to ST sequencing data using the following three steps: 1) Tagging and demultiplexing: the left-most 25 nucleotides (nt) of the R1 sequence contains the 18 nt for the spot-specific barcode and then 7 nt for UMI. The R2 sequence contains the transcript sequence, and its left-most 35 nt are used. The name of every read in the R2 file is tagged with the spot-specific barcode and UMI sequence that are extracted from the paired R1 read. The R2 file is demultiplexed to create the 1007 spot-specific FASTQ files. If the detected spot-specific barcode of a read is not present in the pre-defined barcodes list, the read is excluded from the downstream analysis. 2) Alignment of demultiplexed FASTQ files using Bowtie2^53^. 3) Counting UMI using HTSeq^54^. The reads that are aligned to a user-defined feature (by default: “gene_name”) are collapsed to count only once if they have same UMI. Here we used the version 2.3.1 of Bowtie2^53^ with default parameters and the 0.9.1 version of HTSeq in the “union” mode.

### ST data analysis

UMI counts in each spot were normalized by the total number of transcripts per spot and then multiplied by a scale factor equivalent to the median number of transcripts per spot (TPM). A pseudocount of 1 was added prior to either log10 or Freeman-Tukey transformation. Mitochondrial and ribosomal genes were removed from the expression matrix for further analysis. For PCA of ST data, the 200-700 most variable genes were selected (defined by the Fano factor above a mean-dependent threshold). PC scores for the first six components was then plotted for each spot corresponding to PDAC tissue. For clustering of spots, hierarchical clustering was performed on PC scores using Ward’s criterion on however many components accounted for ≥ 90% of the variance in the data. To extract marker genes specific to each of the resulting region clusters a two-tailed Student’s t-test was used, *P* < 0.01.

### Immunofluorescence staining of FFPE tissue

Immunofluorescent staining of FFPE specimens was carried out using standard procedures. Briefly, sections were cut at 5 µm thickness, and dried overnight. After deparaffinizing the slides, antigen retrieval was carried out by boiling the samples for 10 - 20 minutes in Tris-EDTA or Citrate buffer in a microwave oven. Primary antibodies were diluted 1/100 in Tris Buffered Saline (TBS) + 0.5% BSA and incubated with slides overnight in a wet chamber kept at 4°C. Secondary antibodies (Molecular Probes, Invitrogen) were diluted 1/200 in TBS and incubated with slides for one hour at room temperature prior to mounting and imaging. Fluorescent images were taken on a Hamamatsu NanoZoomer whole-slide fluorescence scanner.

### Determination of cell type enrichment/depletion by multimodal intersection analysis (MIA)

To assess the enrichments of each cell type/subpopulation across tissue regions, marker genes for cell types/subpopulations and ST regions were first identified as described above. We then queried the significance of the overlap between ST genes and cell type marker genes using the hypergeometric cumulative distribution test, with either all genes or the total number of marker genes across all cell types as the background to compute the *P* value. To test for cell type depletion in parallel, if the resulting *P* value was less than 1 - *P*, the −log_10_(1 - *P*) was plotted instead.

## Acknowledgements

We would like to thank C. Loomis, Z. Dewan, B. Dabovic from the NYU Experimental Pathology core, and B. Zeck and L. Chiriboga from the NYU Center for Biospecimen Research and Development (CBRD) for technical assistance, A. Weil from the NYU CBRD for sample acquisition, and members of the Yanai lab for constructive comments.

## Author contributions

R.M. performed the spatial transcriptomics and scRNA-Seq as well as the data analysis. F.W. contributed to scRNA-Seq and spatial transcriptomics analysis. M.C. contributed to spatial transcriptomics and scRNA-Seq processing, and immunofluorescence experiments. M.B. contributed expertise in scRNA-Seq processing and analysis. C.H.H. contributed histology analysis. D.M.S. contributed sample acquisition. I.Y. conceived the project, contributed to the data analysis, and interpretation of the results. RM and IY drafted the manuscript.

## SUPPLEMENTARY TABLES AND FIGURES

**Table S1.** Average number of transcripts per ST spot reported in other publications. The table indicates the number of transcripts detected per spot in other studies using the ST method. Included are only publications or pre-prints that report the average number of detected transcripts.

| Reference | Tissue type | Average UMIs / spot |
| --- | --- | --- |
| Stahl et al Science (2016)<sup>22</sup> | Mouse olfactory bulb | ~30,000 – ~40,000 |
| Asp et al Scientific reports (2017)<sup>26</sup> | Human heart biopsies | ~1,000 – ~3,000 |
| Berglund et al Nat. Comm (2018)<sup>24</sup> | Prostate cancer tissue | 727 – 9019 |
| Maniatis et al Biorxiv (2018)<sup>27</sup> | Human spinal coord samples | 1225 (median) |

**Figure S1.**
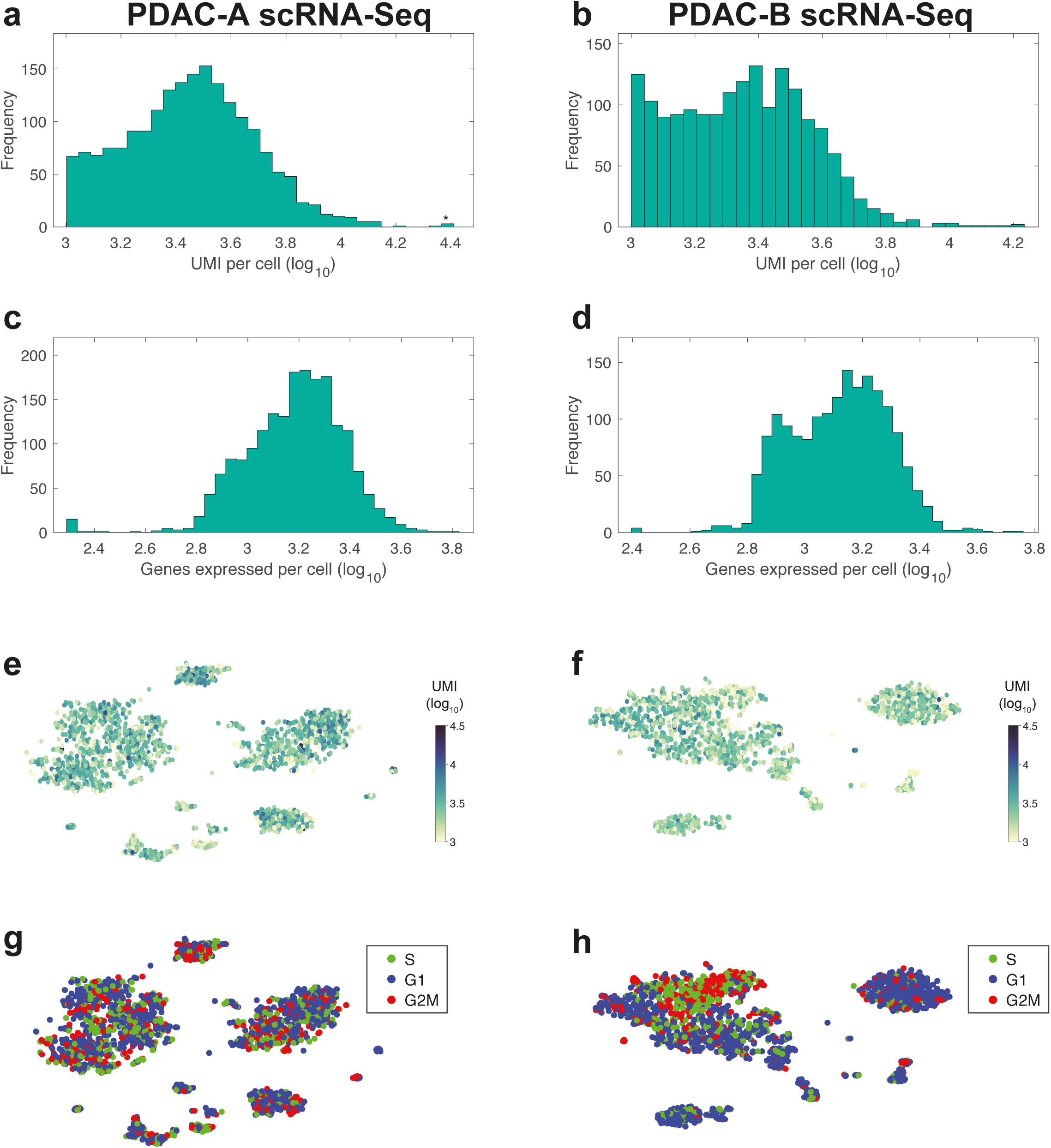
scRNA-Seq data statistics. (a-b) Histogram of unique transcripts detected per cell (log_10_) in PDAC-A (a) and PDAC-B (b). (c-d) Histogram of unique genes expressed per cell (log_10_) in PDAC-A (c) and PDAC-B (d). (e-f) PDAC-A (e) and PDAC-B (f) cells colored by UMIs per cell (log_10_) across tSNE space. (g-h) Cell-cycle phase assignments for PDAC-A (g) and PDAC-B (h) cells mapped across t-SNE space.

**Figure S2.**
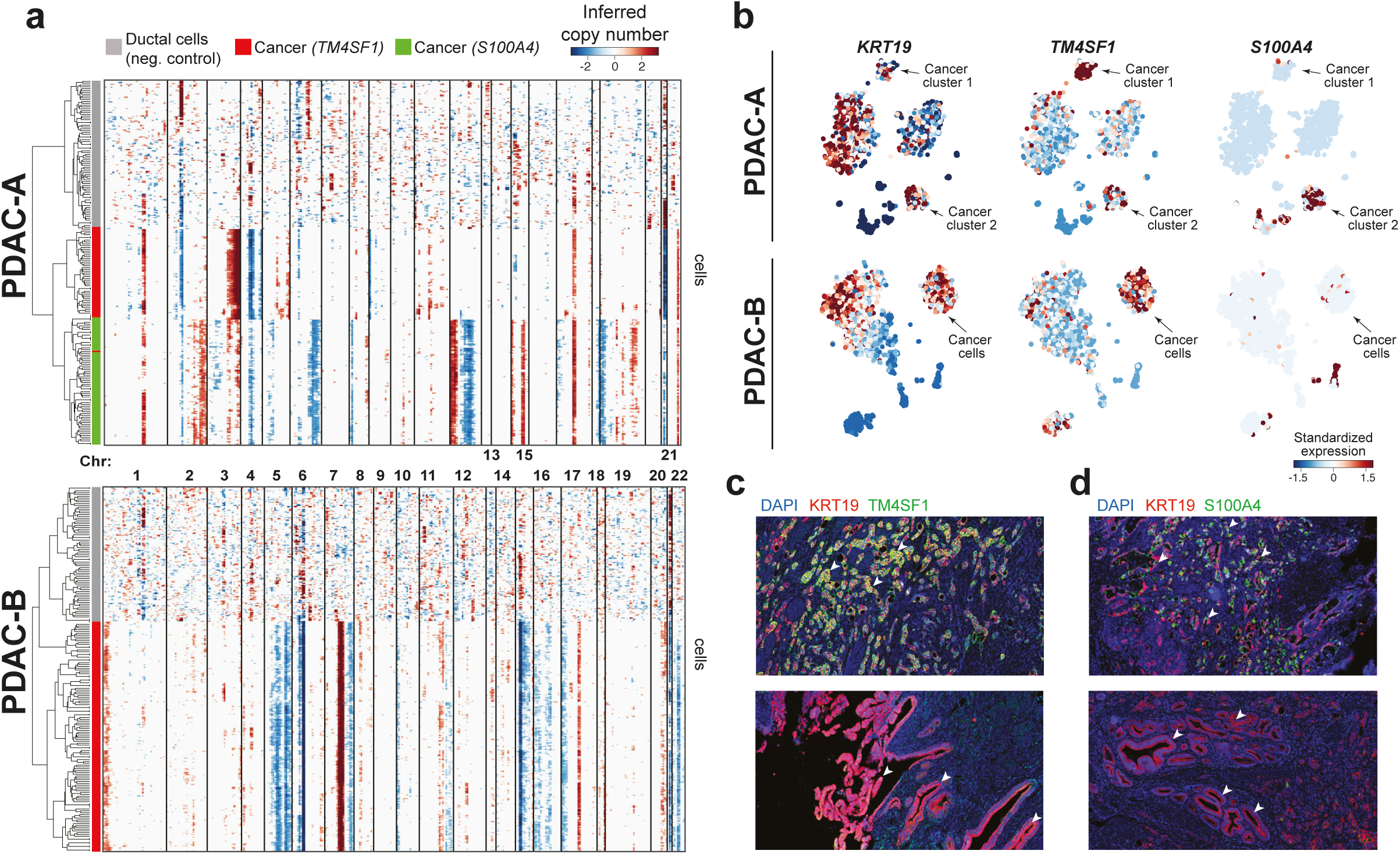
Identification and validation of multiple cancer populations. (a) PDAC-A (top) and PDAC-B (bottom) CNV profiles inferred from scRNA-Seq (same as Fig. 1d,e). A subset of 200 randomly selected ductal cells were chosen as a negative control for the analysis and analyzed together with cancer clusters. Note distance between cancer cells and non-malignant ductal cells in the dendrogram on the left. (b) Expression of *KRT19* (marker of malignant and non-malignant ductal epithelial cells), *TM4SF1,* and *S100A4* projected onto t-SNE of PDAC-A (top) and PDAC-B (bottom). Note specificity of *TM4SF1* expression for PDAC-A cancer A, and *S100A4* expression for PDAC-A cancer B. In PDAC-B, *TM4SF1* is expressed primarily by cancer cells whereas an *S100A4* expressing cancer population is absent. (c) Double immunofluorescence staining for KRT19 and TM4SF1 in PDAC-A FFPE tissue. Note co-localization of KRT19 and TM4SF1 signal in malignant ducts (top panel, white arrowheads), but not in non-malignant ducts (bottom panel, white arrowheads). (d) Double immunofluorescence staining for KRT19 and S100A4 in PDAC-A FFPE tissue. Note co-localization of KRT19 and S100A4 signal in malignant ducts (top panel, white arrowheads), but not in non-malignant ducts (bottom panel, white arrowheads).

**Figure S3.**
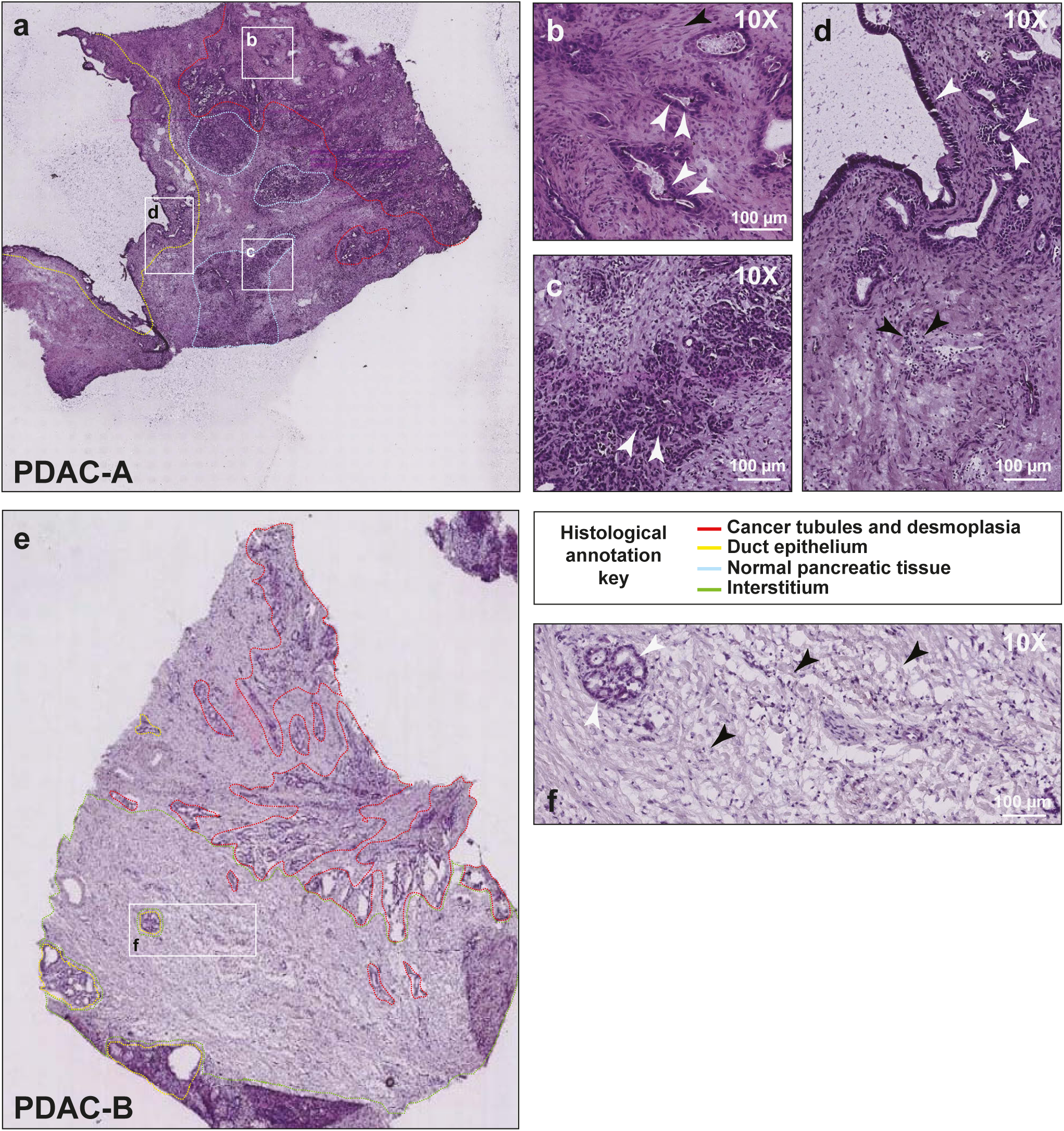
Histology for pancreatic cancer tissue used for spatial transcriptomics. (a) Overview of PDAC-A tissue. (b) Inset of pancreatic cancer ducts and surrounding desmoplasia. White arrowheads indicate cancer cells organizing around tubules. Black arrowheads show the surrounding stroma and desmoplasia. (c) Inset of pancreatic tissue. Arrowheads indicate the acini. (d) Inset of duct epithelium and inflamed tissue. White arrowheads indicate the pancreatic ducts and the black arrowheads point to inflammatory cells with smaller nuclei. (e) Overview of PDAC-B tissue. (f) Inset of PDAC-B tissue showing normal ducts (white arrow) surrounded by interstitial space (black arrows).

**Figure S4.**
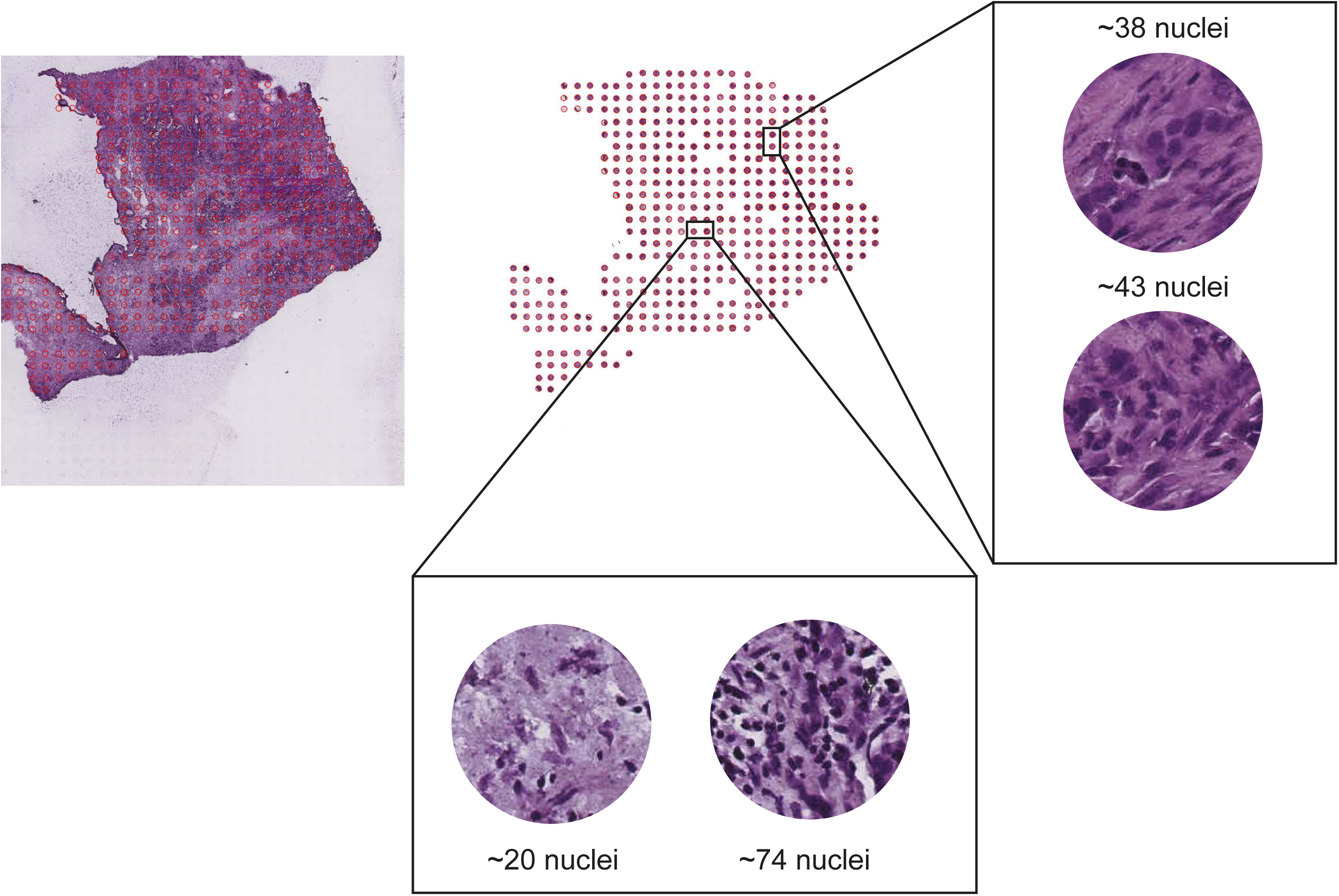
Estimating the total number of cells captured by an ST spot. ST spots were mapped back onto the H&E stained tissue, and brightfield images were extracted from the location of each ST spot. In each enlarged spot, the dark purple dots are nuclei, and the purple background is the cytoplasm and extracellular space. Enlarged spots shown demonstrate that ST spots can capture as few as ten or less cells and as many as a few dozen cells.

**Figure S5.**
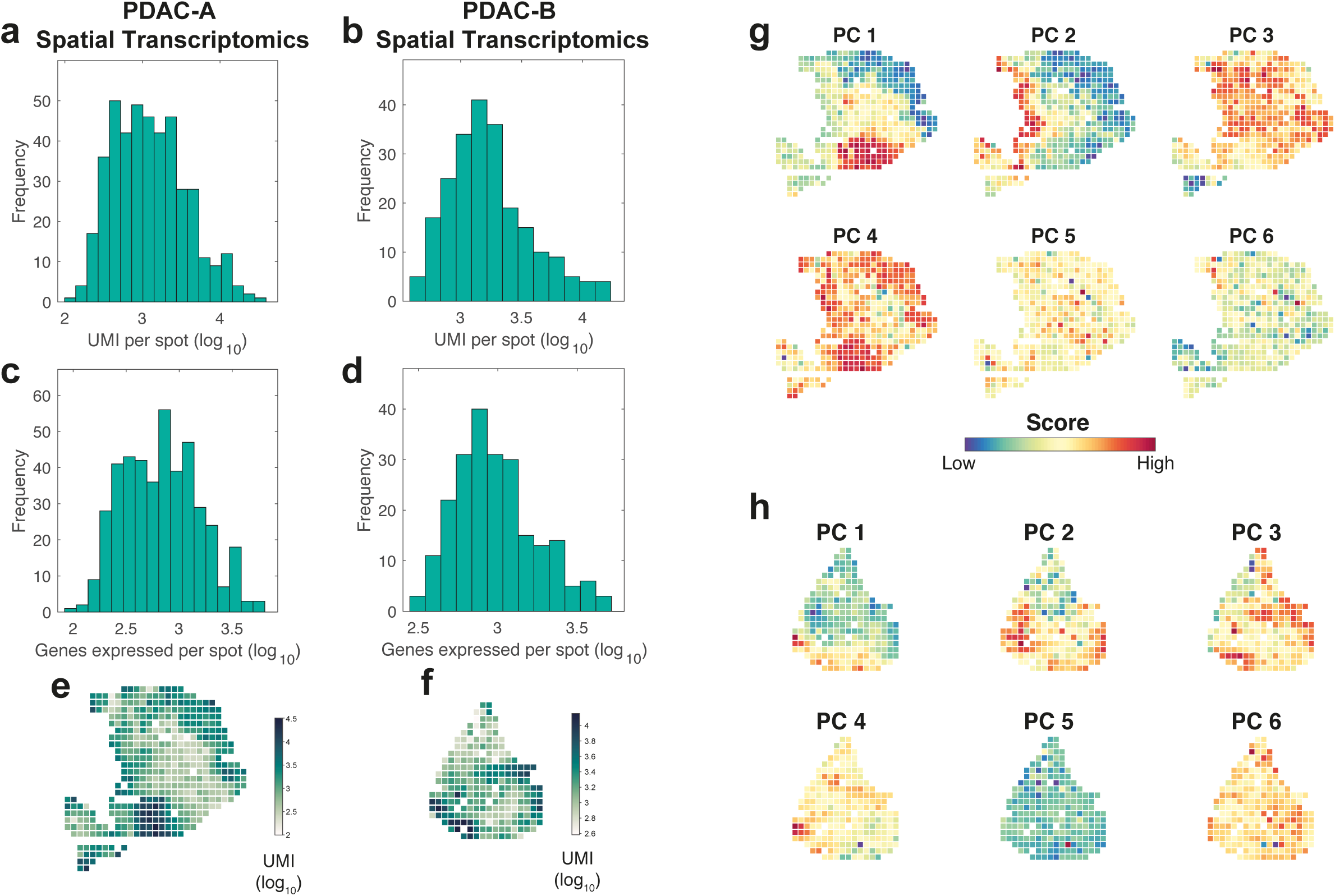
Spatial Transcriptomics (ST) statistics and dimensionality reduction with PCA. (a-b) Histogram of unique transcripts detected per spot (log_10_) in PDAC-A (a) and PDAC-B (b). (c-d) Histogram of unique genes detected per spot (log_10_) in PDAC-A (c) and PDAC-B (d). (e-f) Distribution of unique transcripts detected across ST spots in PDAC-A (e) and PDAC-B (f). (g-h) PC scores for the first six components projected onto ST data for PDAC-A (g) and PDAC-B (h).

**Figure S6.**
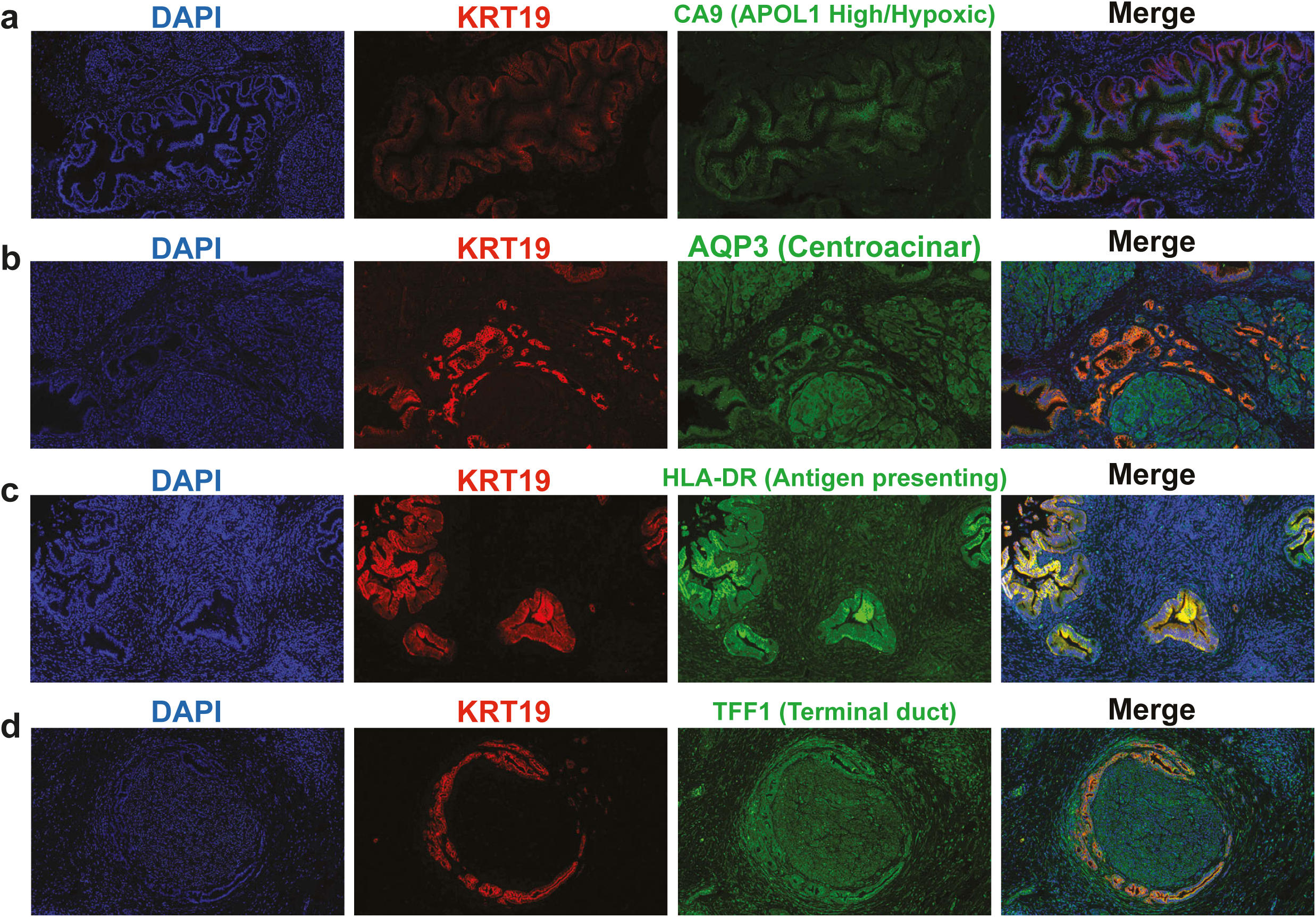
Immunofluorescence staining of ductal subpopulation markers in PDAC tissue. FFPE tissue was co-stained for KRT19 (duct marker) and subpopulation markers CA9 (a), AQP3 (b), HLA-DR (c), and TFF1 (d), as shown in Figure 3. Individual signals are shown here in addition to the merged signal to better appreciate marker co-localization.

**Figure S7.**
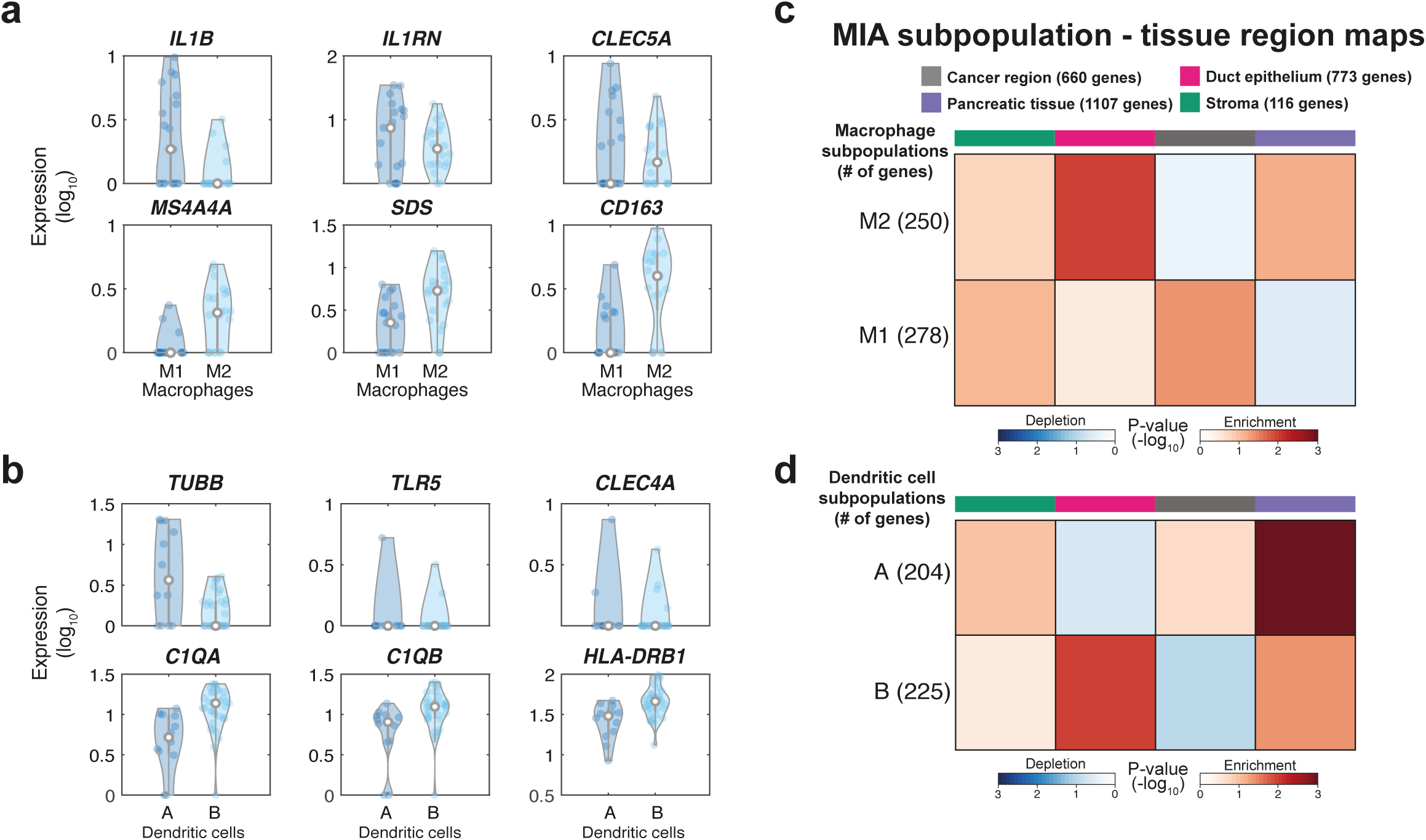
MIA maps of macrophage and dendritic cell subpopulations across PDAC-A tissue. (a) Expression of M1 (top row) and M2 (bottom row) marker genes (two-tailed Student’s t-test, *P <* 10^-5^). (b) Expression of dendritic cells A (top panel) or B (bottom panel) marker genes (two-tailed Student’s t-test, *P <* 10^-5^). (c-d) Enrichment of macrophage (c) and dendritic cell (d) subpopulations across PDAC-A ST regions. Indicated are the number of genes used for MIA.

